# Robust gene expression-based classification of cancers without normalization

**DOI:** 10.1101/2020.04.28.051953

**Authors:** Aixiang Jiang, Laura K. Hilton, Jeffrey Tang, Christopher K. Rushton, Bruno M. Grande, David W. Scott, Ryan D. Morin

## Abstract

Binary classification using gene expression data is commonly used to stratify cancers into molecular subgroups that may have distinct prognoses and therapeutic options. A limitation of many such methods is the requirement for comparable training and testing data sets. Here, we describe and demonstrate a **s**elf-**t**raining implementation of **p**robability **r**atio-based classification **p**rediction **s**core (PRPS-ST) that facilitates the porting of existing classification models to other gene expression data sets. We demonstrate its robustness through application to two binary classification problems in diffuse large B-cell lymphoma using a diverse variety of gene expression data types and normalization methods.

## Background

The classification of tumors into molecular subgroups using gene expression features has been applied to numerous cancer types and can be used to identify high-risk patients and/or determine suitable treatment options [1–4]. Accordingly, clinical-grade assays using such classifications can serve as robust prognostic or predictive biomarkers and in some cases aid in obtaining a molecular diagnosis for neoplasms that are histologically indistinguishable. The implementation of clinical assays, however, depends on the ability to accurately and reproducibly classify tumors in the research setting such that shared biological or genetic features underlying these subgroups can be characterized.

A variety of machine learning or modeling approaches have been applied to the binary classification of cancers, such as support vector machines (SVM) [5], penalized regression [6,7], and k nearest neighbor clustering [8,9], to name a few. However, all of these methods require comparable training and testing data sets. Here, by comparable we mean that both data sets are sufficiently similar that they can be considered unbiased, random samples from the same population. As truly comparable data is rarely available due to inter-experimental variation and platform differences, expression data from unlabeled cases (testing cohorts) are commonly normalized to an available training cohort. Such normalization has the potential to remove signal and increase noise and can be intractable when working with data from distinct platforms [10,11]. Platform differences can also lead to incomplete matching of variables between data sets, for example when data are generated using different microarray designs or divergent protocols for RNA-seq library handling or post-processing. In these situations, classifiers can only be ported by re-modelling on the training data set using shared variables, which leads to differences in their coefficients (or weights).

An established classification system in diffuse large B-cell lymphoma (DLBCL) is termed “cell of origin” (COO), in which the activated B-cell-like (ABC) subtype is associated with inferior outcomes relative to the germinal centre B-cell-like (GCB) subtype, and cases are considered unclassified (U) when they cannot be confidently assigned to either group with an empirical Bayes probability cutoff of 0.9 [12–14] A further extension to COO was recently established that identifies predominantly GCB-DLBCL tumors expressing the double-hit gene expression signature (DHITsig), which also identifies a group of patients with inferior outcomes [15]. The conventional application of COO to separate ABC and GCB tumors relies on the linear predictor score (LPS) method, in which weights are obtained for genes that have been established as up- or down-regulated in one of the two subgroups using a training cohort [13]. Using these weights, a LPS can be obtained from additional cohorts that have been normalized to the training data [13]. The gene-wise normalization step assumes that the distribution of expression of each gene and the proportion of tumors of each subtype are roughly the same in both training and testing data sets, which may not be a valid assumption due to sample selection bias and variable representation of molecular subgroups in different populations. The Lymph2Cx and DLBCL90 NanoString assays apply the LPS method to a smaller number of fixed genes and uses housekeeping genes for normalization [15,16]. Importantly, when Lymph2Cx has a new codeset, a set of standards with known LPS scores are processed and linear regression is used to calibrate LPS scores for the new codeset. Although this is an effective approach for a robust clinical assay, it is not applicable in the context of discovery-based gene expression experiments.

To eliminate the constraints associated with existing classification methods, we have extended our previously described binary classification method “**p**robability **r**atio-based classification **p**rediction **s**core” (PRPS; pronounced “porpoise”) to allow self-training of cases in each subgroup without any labels (PRPS-ST) [15]. The algorithm requires a set of weights for genes that distinguish the two classes, which can be derived from any gene expression dataset with high-confidence class labels. Importantly, we show that weights derived from RNA-seq data can be applied to accurately classify samples with RNA-seq data analyzed through disparate alignment and quantification pipelines. More strikingly, we show that PRPS self-training facilitates robust porting of classifiers to data from hybridization-based gene expression platforms including microarray and Illumina DASL. While we have focused here on the application of the method to DLBCL classification, we expect this method could prove useful in any binary classification based on gene expression differences.

## Methods

### Self-training approach for PRPS

The goal of the self-training algorithm is to identify cases in an unlabeled gene expression data set that can be confidently classified (“Self-training”, Figure 1), and to use those cases as pseudo training data thereby allowing classification of all cases in the cohort (“Empirical Bayes Classification”, Figure 1). Besides a gene expression matrix, the algorithm requires a set of *m* genes with a weight *w*_*k*_ for each gene *k*.

**Figure 1:**
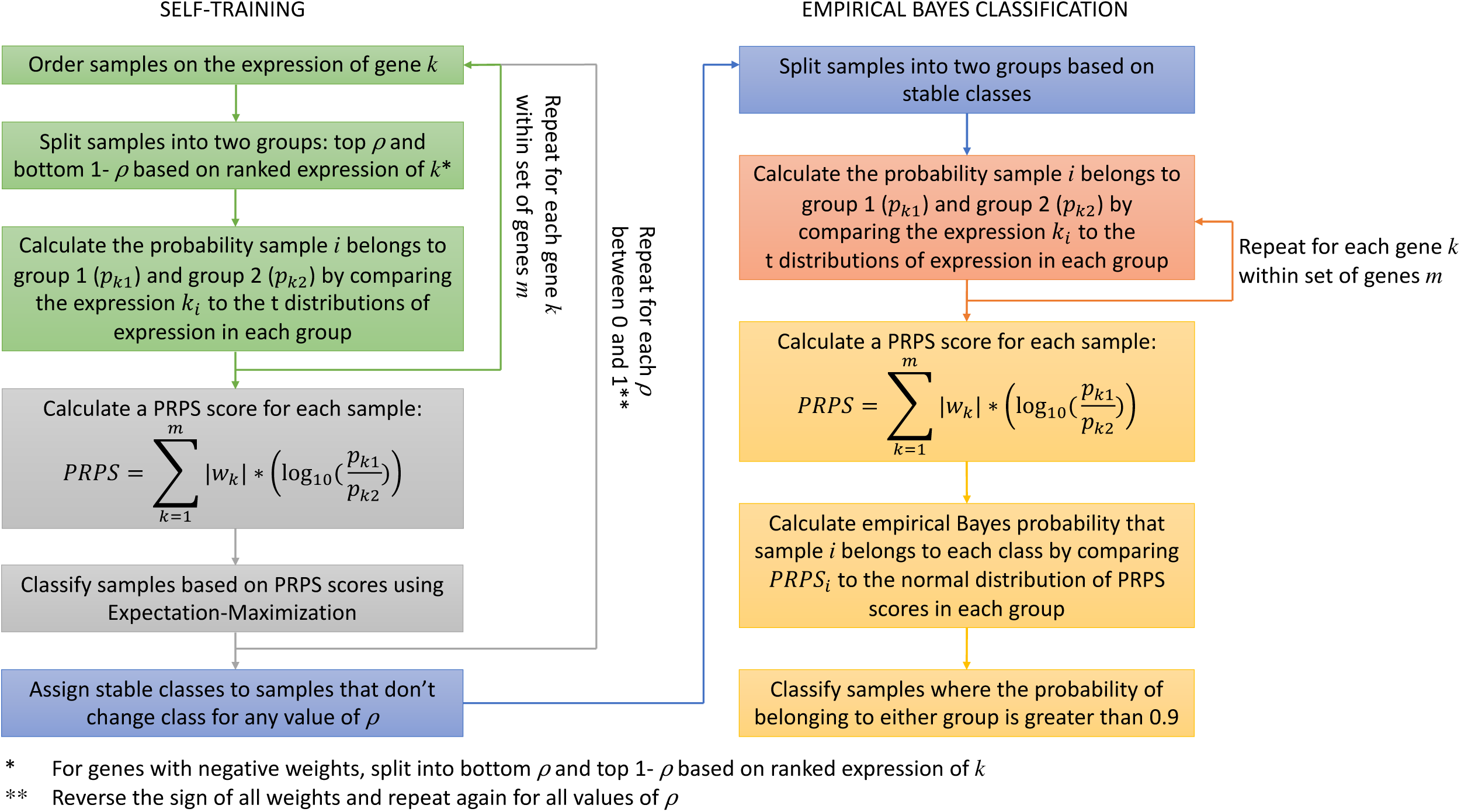
Flow chart of self-training algorithm illustrated by PRPS classification. The self-training process (left) searches for stable classes; Empirical Bayes classification (right) performs binary classification based on stable classes.

During the self-training stage, we iteratively divide tumors into two classes, enforcing the relative proportion of each class as *ρ* and 1-*ρ*. This is performed over a range of *ρ* values spanning (0,1) individually for each gene. Here, we use the package default search range for *ρ*, 0.05-0.95, at intervals of 0.05. For each gene *k* with a positive weight *w*_*k*_ the samples are split into two groups representing tumors with the highest *ρ* and lowest 1 − *ρ* expression of *k*. If *w*_*k*_ is negative, the tumors are split according to the lowest *ρ* and highest 1 − *ρ* expression of *k*. Next, stabilized *t* values *t*_*ik*1_ and *t*_*ik*2_ are calculated for each tumor *i* as follows:

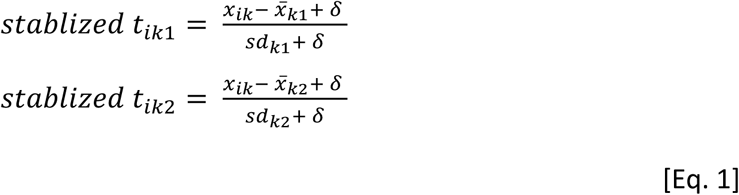

Here, *x*_*ik*_ is the expression of gene *k* for the *i*^*th*^ tumor, and 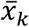 and *sd*_*k*_ are the mean and standard deviation of expression of gene *k* within each group. We stabilize the *t* values to avoid inflation and breaking up of continuity by adding a small number (by default, 0.01), *δ*, to both the numerator and denominator. We then estimate the probability that sample *i* belongs to group 1 (*p*_*ik*1_) or group 2 (*p*_*ik*2_) by comparing *t*_*ik*1_ and *t*_*ik*2_ against *t* distributions with degrees of freedom determined by the sample size of each group, respectively. These steps are repeated until all *p*_*ik*1_and *p*_*ik*2_have been calculated. A PRPS score is then calculated for each tumor as follows:

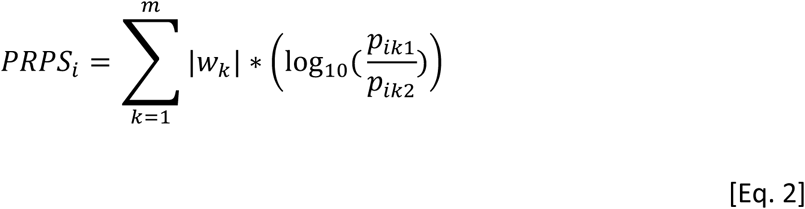

The PRPS scores are used to classify samples into two groups using Expectation Maximization (EM), performed with the mclust R package version 5.4.5 [17]. All of the above steps are repeated for each value of permutated *ρ* spanning (0,1). To increase variation, this process is also repeated with the sign of all weights reversed. The samples that have the same classification for all values of *ρ* with original weights and sign-reversed weights are assigned a stable class label.

During the Empirical Bayes classification portion, these stable class labels derived from self-training portion are used to split the samples once again into two groups, and *t*_*ik*_ and *p*_*ik*_ are calculated for each group using the same method as in the self-training portion of the algorithm. PRPS scores are again calculated for each tumor using Eq. 2. Lastly, we calculate an empirical Bayes probability that a sample belongs to one of the two groups as follows:

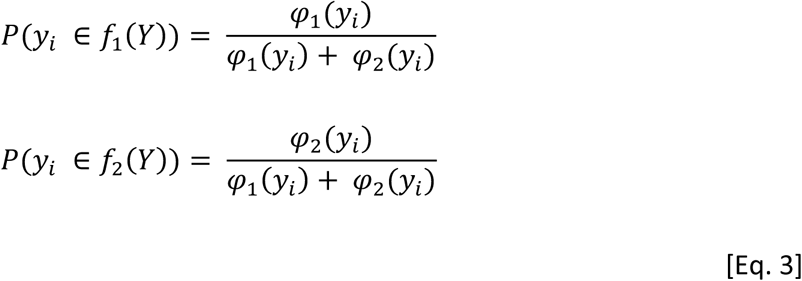

Here, *φ*_1_(*y*_*i*_) is the estimated density value of the *i*^*th*^ sample under the normal distribution 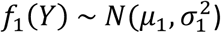 of PRPS scores, where the mean and standard deviation (SD) are estimated from samples in the group 1 stable class. Similarly, *φ*_2_(*y*_*i*_) is the estimated density value of the *i*^*th*^ sample under the normal distribution 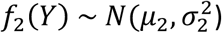 of PRPS scores. A probability threshold of 0.9 is set for inclusion in either classification, and samples with probabilities below this threshold for both groups are unclassified. This step is identical to the estimation of probabilities from LPS scores [13]. Self-training functions are included in the PRPS R package, which is available from CRAN and on GitHub (https://github.com/ajiangsfu/prps).

### Obtaining feature weights

The self-training algorithm requires a set of feature weights for genes whose expression levels vary significantly between groups. We previously described a method to obtain weights for DHITsig classification by taking the mean of four scaled importance scores and we have used those weights here for the DHITsig binary classifications [15]. Weights for COO classification were generated using the Ennishi *et al*. RNAseq data for which Lymph2Cx-generated COO labels are available [15]. Briefly, the RNAseq reads were aligned with STAR and quantified using featureCounts followed by variance stabilizing transformation (VST) in DeSeq2 [18–20]. The weights are the *t* values from a *t* test comparing the expression of each gene between the ABC and GCB cases as determined by Lymph2Cx. This is comparable to how weights were derived for the original LPS COO classification method [13]. We generated weights for each of 153 Wright genes as used by Morin et al., 2011 [21] and separately selected 100 genes with differential expression between the two COO groups based on smallest P value (“top 100”) for use in COO classification. All weights are provided in Supplemental Tables S1 and S2.

### Gene expression and mutation data

Our objective in selecting cohorts was to demonstrate the transferability of the self-training algorithm to many different types of gene expression data using cancers with established binary classification systems. We used gene expression data from Reddy et al., Schmitz et al., and Scott et al. to test the accuracy of the PRPS self-training for COO classification, and the REMoDL-B data for DHITsig classification [16,22–24]. Details of each data set are in Table 1.

**Table 1.**
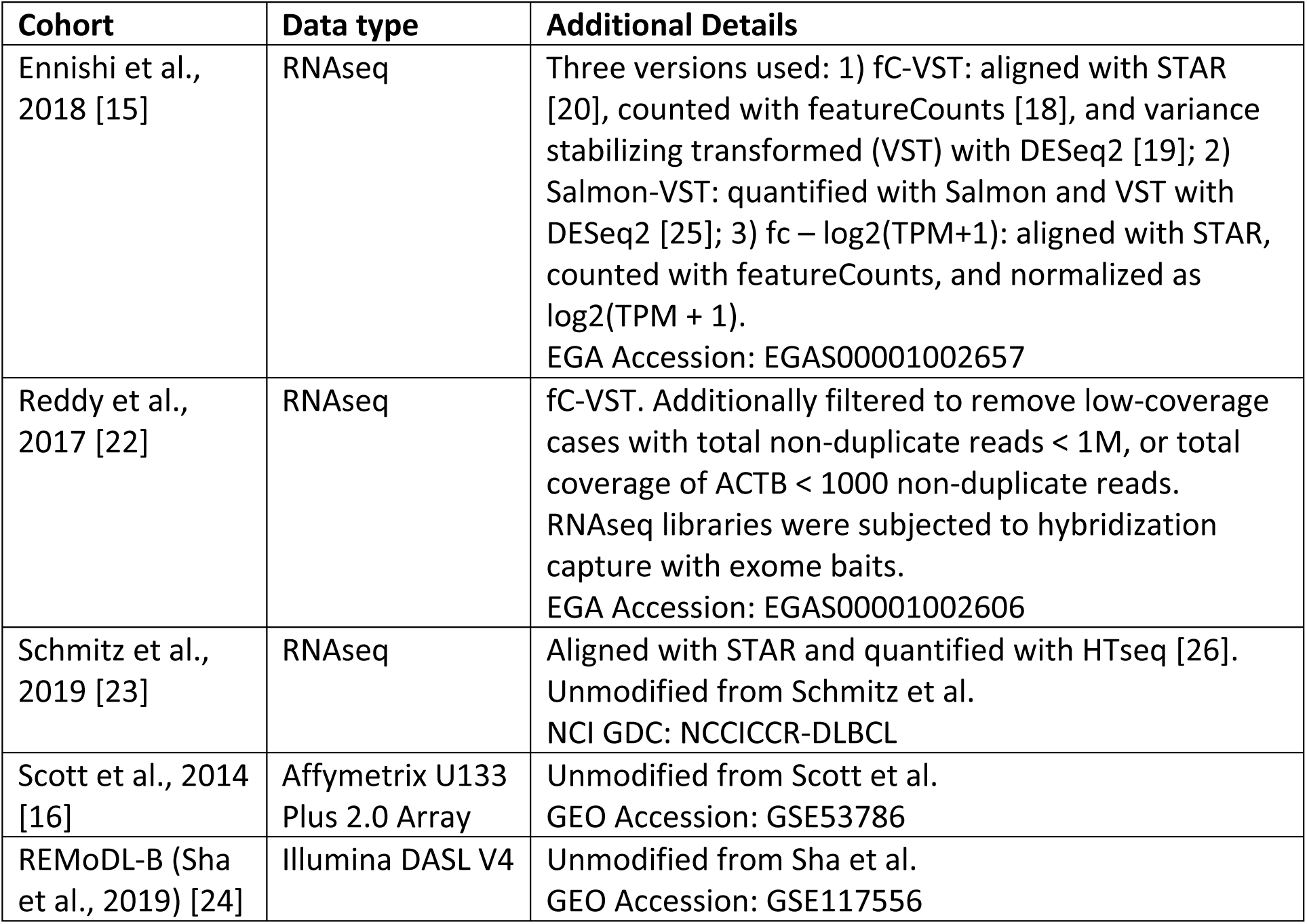
Details of gene expression data used to test the PRPS self-training classification.

Exome sequencing data from Reddy *et al*. and Schmitz *et al*. were obtained from the European Genome-Phenome Archive (Reddy: EGAS00001002606) and National Cancer Institute Genomic Data Commons (Schmitz: NCICCR-DLBCL). Using the original alignments, data were reanalyzed using a standardized variant calling pipeline for tumor samples with no matched constitutional DNA. Candidate simple somatic variants were identified using Strelka2 [27], utilizing small insertions and deletions identified using Manta [28]. Candidate variant positions were then converted to BED format and provided to GATK4 MuTect2 (https://gatk.broadinstitute.org/hc/en-us/articles/360036730411-Mutect2), leveraging a panel of normals generated from 58 unrelated normal genomic samples. Variants were annotated using vcf2maf (https://github.com/mskcc/vcf2maf), and any variants with a population allele frequency of >0.005 in gnomAD [29] were flagged as germline and removed. Variant calls were further post-filtered to remove those with 1) Less than 5 reads supporting the alternate allele, 2) Read mean mapping quality < 50, 3) Read mapping strand bias p<0.01, determined using a Fisher’s exact test between the reference and alternate allele, 4) Base quality bias p<0.01, determined using a t-test on all bases at a variant position, and 5) variant allele frequency >0.01. Finally, variant calls generated from the Schmitz cohort were lifted over to hg19 using Crossmap [30]. *BCL2* and *MYC* break apart fluorescence *in situ* hybridization (FISH) data were available for REMoDL-B.

### PRPS self-training and COO and DHITsig classification

For COO classification in each cohort, expression matrices were subset to include only matched Wright or Top 100 genes (Supplemental Tables S1 and S2). In the Reddy cohort, many of the top 100 weighted genes had very low read counts. We attribute this to the hybridization capture methodology applied to RNAseq libraries prior to sequencing. To address this, we considered genes with low expression as missing data and included only the 71 genes having a mean variance stabilized expression value > 6. In the Scott cohort, only 69/100 genes matched, and 68/100 matched in the Schmitz cohort. At least 150 of the Wright genes were matched in each cohort. All genes used for COO classification, along with their weights and their representation in each cohort are shown in Supplemental Table S1. For DHITsig classification, 97/104 DHITsig genes with weights from Ennishi et al. were represented in the Illumina DASL data of the REMoDL-B cohort (Supplemental Table S2). PRPS was run on each expression matrix using default parameters (ratio search range 0.05-0.95, probability cutoff of 0.9).

### Evaluating the effect of sample size on accuracy of self-training

To determine the lower cohort size limit for self-training, we applied a random sampling procedure using the Scott and Reddy cohorts for COO and REMoDL-B for DHITsig. The maximum number of cases in each sample was determined by cohort size. For the Scott cohort, our maximum total sampling sample size was 110 and we arbitrarily chose a minimum total sampling sample size of 20. For consistency, we used the same range of sample sizes for the Reddy cohort. Using steps of 10, we sampled across the range of sample sizes *m* = 20, 30, …, 110. For each given sample size *m*, we randomly sampled *m* cases from a given cohort without any repeated cases. We repeated this procedure for 1000 times for each given sample size.

We used the REMoDL-B cohort to evaluate the accuracy of DHITsig classification, which naturally has a minority class (DHITsig+). We preserved the class ratios by sampling proportionately from DHITsig+ and DHITsig-GCB cases. Again, all samplings were random and without replacement. We repeated this procedure 1000 times for each given sample size.

### Identifying COO-enriched mutations

Mutations from a targeted sequencing a panel of lymphoma-related genes [31] were reduced to a mutation matrix where known targets of non-synonymous or hotspot mutations were binary coded as mutated or unmutated. Known targets of aberrant somatic hypermutation (aSHM) were coded according to the number of mutations within the typical target region for aSHM, defined for each gene as the region proximal to the transcription start site containing high frequency of either coding and non-coding mutations. The matrix was filtered to include only genes, hotspots, or aSHM regions that were mutated in at least 10% of tumors. We used Fisher’s exact tests to identify features significantly enriched in either subgroup using a Benjamini-Hochberg correction and a false discovery rate (FDR) threshold of 0.1. From these data, we identified the six loci most enriched for mutations in the ABC subgroup (MYD88 Codon 273, ETV6 Any, PIM1 Hypermutated, NFKBIZ 3’ UTR, TMSB4X Nonsynonymous, IRF4 Nonsynonymous) and five loci most enriched for mutations in the GCB subgroup (GNA13 Nonsynonymous, TNFRSF14 Nonsynonymous, BCL2 Hypermutated, EZH2 Codon 646, P2RY8 Nonsynonymous), while excluding regions with insufficient coverage in the Reddy and Schmitz exome sequencing data.

To allow comparison with the COO labels from PRPS, the Reddy and Schmitz cases were assigned as either ABC- or GCB-mutated if they were mutated in one or more of the defined regions. For regions known to be affected by aSHM, a case was considered mutated if it had >2 mutations within the defined aSHM region regardless of the effect of the mutation on protein. McNemar’s tests were then used to compare the number of ABC- or GCB-mutated cases that were correctly classified by each subgrouping method.

### Statistical Analysis

The Kaplan-Meier method was used to estimate progression-free survival (PFS) or overall survival (OS) within different COO or DHITsig classifications. Univariable and multivariable Cox proportional hazard models were used to evaluate and compare different classification methods. PRPS scores were tested for correlation with other scores using Pearson correlation. Two-class accuracy was calculated for samples that were classified (ABC/GCB or DHITsig+/-) by both methods being compared, while three-class accuracy included cases called “unclassified” by either method. Two-class accuracy was also calculated for samples that were classified for DHITsig+ vs others for sample size sampling experiments. A threshold for significance of *P* < 0.05 was employed for all tests. Where appropriate, Benjamini-Hochberg multiple test correction was applied with a FDR threshold of 0.1. All statistical tests were performed in R-3.5.

## Results

### Self-training COO classification accuracy on digital and analog gene expression data

We used our cohort of 272 DLBCLs with Lymph2Cx-determined COO labels from Ennishi et al., 2019 [9] to obtain *t* values for 155 COO-distinguishing genes used in previous LPS-based classification models, herein referred to as the Wright genes. As these genes were selected using microarray data and are not necessarily ideal for RNA-seq, we separately identified the top 100 differentially expressed genes and used the *t* values of either gene set as weights. 45 of the top 100 differentially expressed genes were shared with the Wright list and, notably, many of those unique to the latter had small *t* values both in our cohort and the Reddy cohort, which has the potential to introduce noise in the classification (Figure S1).

We compared the self-training algorithm on the Ennishi RNAseq data quantified using three commonly used read-counting methods, namely variance stabilized gene-level read counts from Salmon (Salmon-VST), log-transformed TPMs inferred by featureCounts (fc-log2(TPM+1)), and variance stabilized featureCounts data (fC-VST). The PRPS scores determined using all three data formats were highly correlated (Figure S2A). Accordingly, each data processing method yielded similar two-class accuracy (ABC and GCB), with all exceeding 0.97 (Supplemental Table S3). Three-class accuracy (ABC, GCB, and U) was generally lower due to variation in the number of cases deemed unclassifiable. Overall, this demonstrates the consistency of the self-training tool on RNA-seq data processed through different pipelines (Figure S2B and C).

We next validated the self-training algorithm using the microarray data from Scott et al. [16]. In addition to Lymph2Cx calls, the Scott cohort has COO calls generated with the original LPS algorithm and is considered the “gold standard” [12]. Using expression data for the 151 matched Wright genes and the 69 matched Top 100 genes we classified cases using the PRPS self-training algorithm and used heat maps to visualize expression patterns between subgroups (Figures 2A and S3A). Compared to the gold standard class labels, the self-training classifications with either the Top 100 (2 class accuracy: 98.95%, 3 class accuracy: 85.71%) or Wright gene weights (2 class accuracy: 100%, 3 class accuracy: 83.19%) were as accurate as Lymph2Cx (2 class accuracy: 98.86%, 3 class accuracy: 82.76%; Supplemental Table S3). Moreover, the PRPS scores determined by the self-training algorithm were strongly correlated with both gold standard and Lymph2Cx scores (Figure 2B-D, S3B-D). A high hazard ratio is maintained for the ABC class compared to the GCB class when these newly classified cases are considered, suggesting they have been classified appropriately (Figure 2E, Table 2).

**Table 2.**
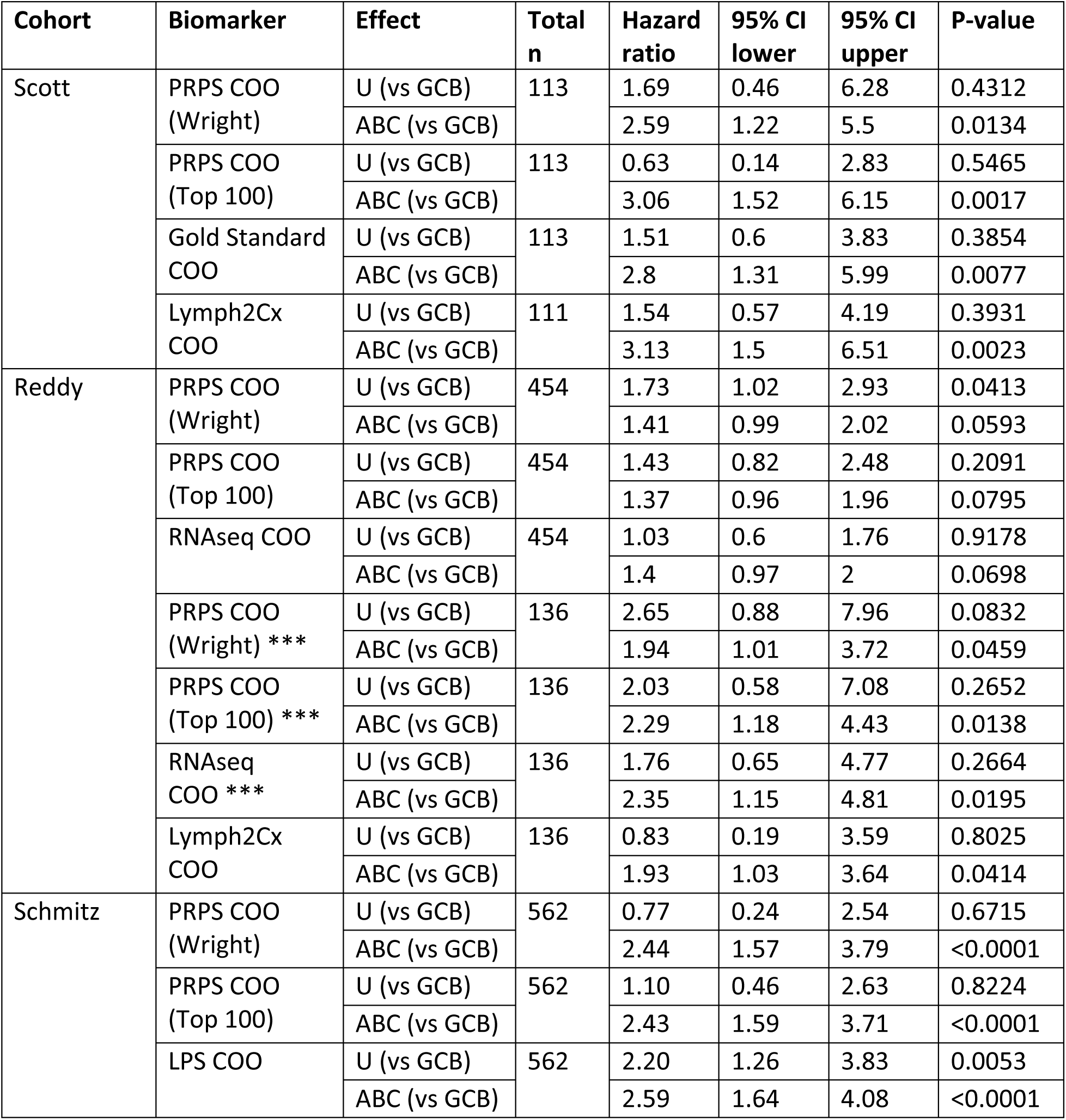
Cox models comparing overall survival between COO subgroups as determined by different methods. ***Cox models performed using only cases with Lymph2Cx labels.

**Figure 2:**
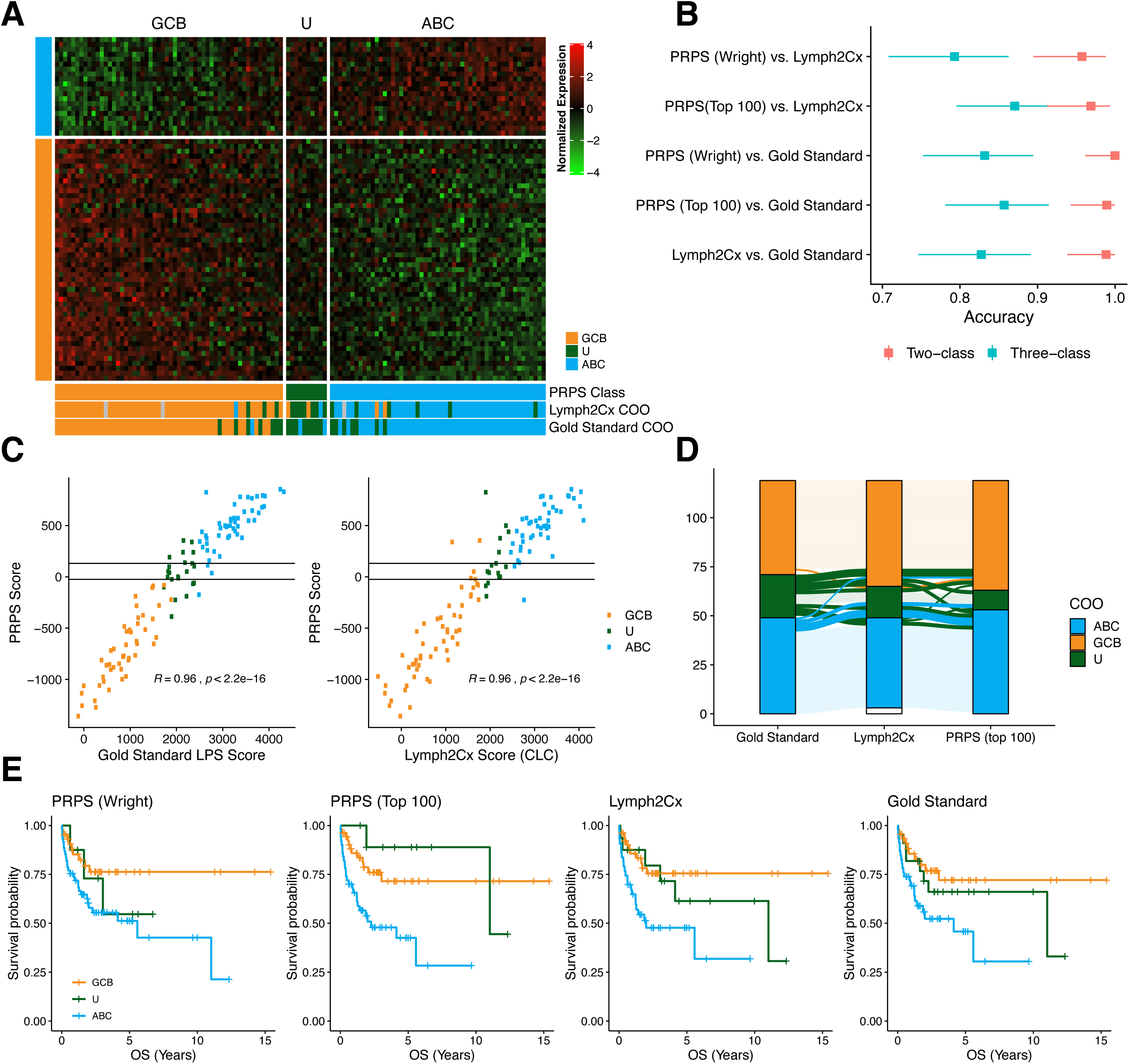
Comparing the performance of self-training PRPS COO classification to gold standard LPS and Lymph2Cx on the Scott cohort. **A**. Heatmap of 69 matched “top 100” COO genes. Samples are ordered by PRPS score. Expression data are normalized per gene to a mean of 0 and standard deviation of 1. **B**. Two- and three-class accuracy comparisons between classification methods. Error bars represent 95% confidence interval. **C**. Scatter plots comparing scores derived from different classification methods. R and *p* values from Pearson correlation. **D**. An alluvial plot reveals how cases move between classes with different classification methods. The lines between classifications are colored according to their gold standard classification. **E**. Kaplan-Meier OS survival curves for each COO classification method.

In the study by Reddy et al., COO labels were derived from their RNA-seq data using a bespoke algorithm based on the difference of mean standardized expression values of 11 ABC and 9 GCB genes (“RNAseq ABC/GCB”). The Lymph2Cx assay was also applied 137/468 cases. Using Lymph2Cx classifications as ground truth, our self-training PRPS classifications were more accurate (Top 100 genes: 2 class accuracy: 97.48%, 3 class accuracy: 86.13%; Wright genes: 2 class accuracy: 96.67%, 3 class accuracy: 88.32%) than the RNAseq ABC/GCB (2 class accuracy: 95.58%, 3 class accuracy: 82.48%) (Figure S4A, Supplemental Tables S16). PRPS scores correlate strongly with the RNAseq ABC/GCB and Lymph2Cx score with either the top 100 (Figure S4B) or Wright gene weights (Figure S4C).

The Schmitz data had COO labels generated using the original LPS method, but LPS scores were not provided and the Lymph2Cx assay was not used. To allow comparison of our self-training results to LPS, we used our implementation of the LPS method with the Wright genes to generate LPS scores, which issued 100% 2 class accuracy and 93.95% 3 class accuracy (Supplemental Table S3). Two factors may account for the small difference in accuracy: first, the Schmitz LPS method used 195 genes to our 153; and second, they performed post-processing with adjusted weight normalization to their previous microarray U133+ data before LPS classification. Scores from our in-house LPS and self-training methods correlate strongly when either gene set is used for self-training classification (Figure S5A). Using the LPS COO labels assigned to these data as truth, the self-training COO labels generated with either gene set yielded very similar two-class accuracy but the latter gene list had inferior three-class accuracy (Figure S5B).

### Assessing accuracy using subgroup-restricted driver mutations

In all of the cohorts described above, PRPS self-training COO classification had a very low frank misclassification rate, but the unclassified (U) group was consistently smaller relative to the other methods. To objectively assess whether this results from true ABC and GCB DLBCLs becoming correctly classified by PRPS, we examined whether tumor genetic features were consistent with their class assignment in the cohorts with available mutation data. Although many genes have been reported as more commonly mutated in either ABC or GCB DLBCL, the strength of these associations varies by gene. Using the targeted sequencing data from Ennishi et al., we first identified the 6 ABC and 5 GCB features most strongly enriched for mutations, respectively, based on Lymph2Cx COO labels and used these for subsequent analyses (Figure S6).

Using this gene list, the cases in both the Reddy and Schmitz cohorts were identified as either ABC- or GCB-mutated if each had at least one COO-characteristic mutation. In the Reddy cohort, self-training with the top 100 genes correctly classified significantly more GCB-mutated tumors than the RNAseq ABC/GCB method (McNemar’s test), while there was no significant difference in the number of correctly classified ABC-mutated tumors (Figure 3A, Table 3). Surprisingly, although the prognostic difference between patients stratified on COO is well established, none of the classification methods resulted in a significant survival difference between ABC and GCB in the Reddy cohort when all cases were included (Figure 3B, Table 2). In order to directly compare our classifications to Lymph2Cx, we also generated Cox models for the subset of cases with Lymph2Cx labels. The PRPS self-training classifications produced a HR as great as or greater than that obtained with the Lymph2Cx classification (Figure 3B). In the Schmitz cohort, PRPS self-training classification with the top 100 genes was not significantly more or less accurate than the original LPS COO classification (Table 3). However, self-training with the Wright genes accurately classified significantly more ABC-mutated tumors but significantly fewer GCB-mutated tumors than LPS (Figure 3C, Table 3), demonstrating that the Wright genes may lead to under-calling GCB tumors with self-training classification. All of the COO classifications on the Schmitz data maintained significant survival differences with a high hazard ratio associated with the ABC subtype relative to GCB (Figure 3D, Figure S5C, Table 2).

**Table 3.**
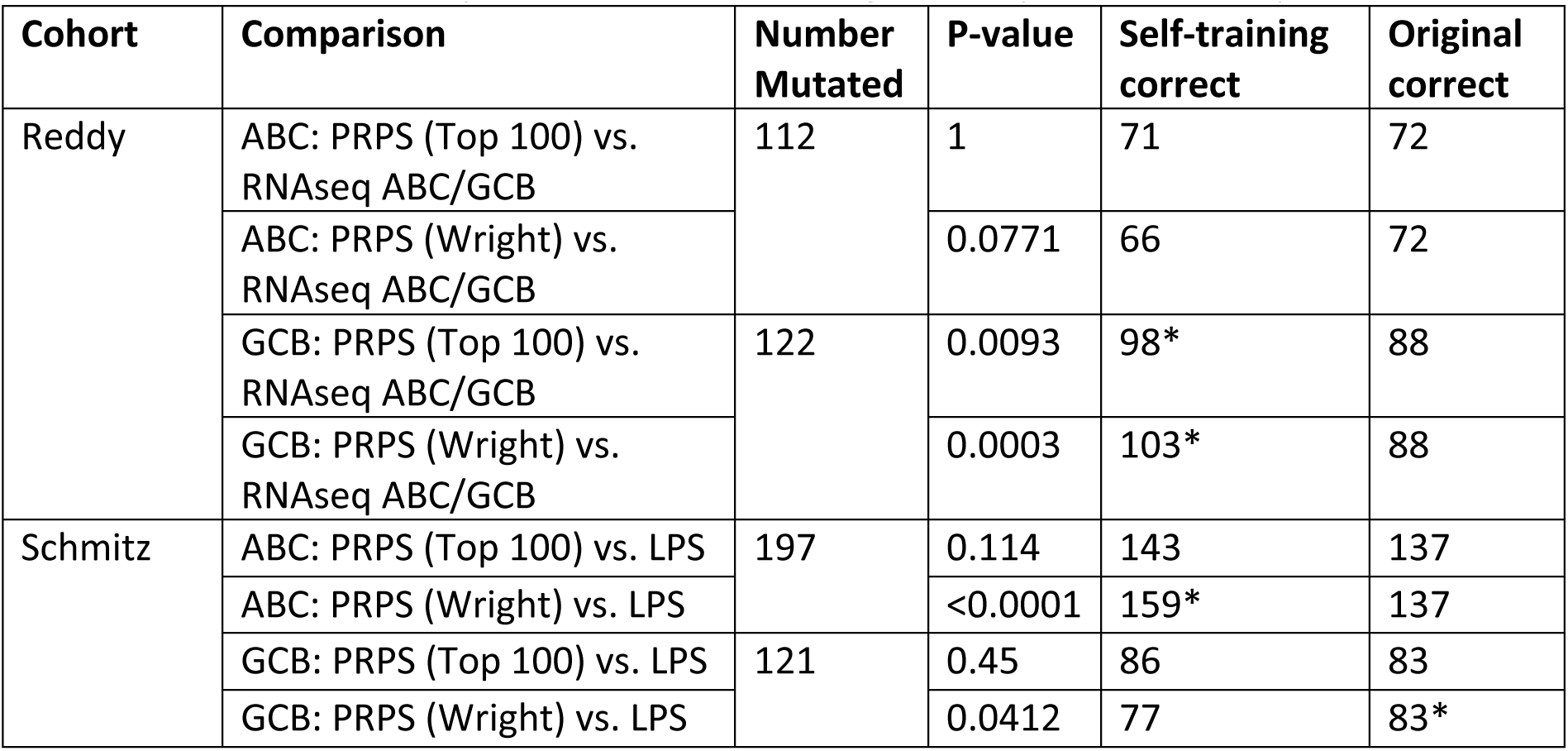
McNemar’s Tests comparing the number of correctly-classified cases based on mutation status achieved by different methods. *Significantly more correctly-classified cases.

**Figure 3:**
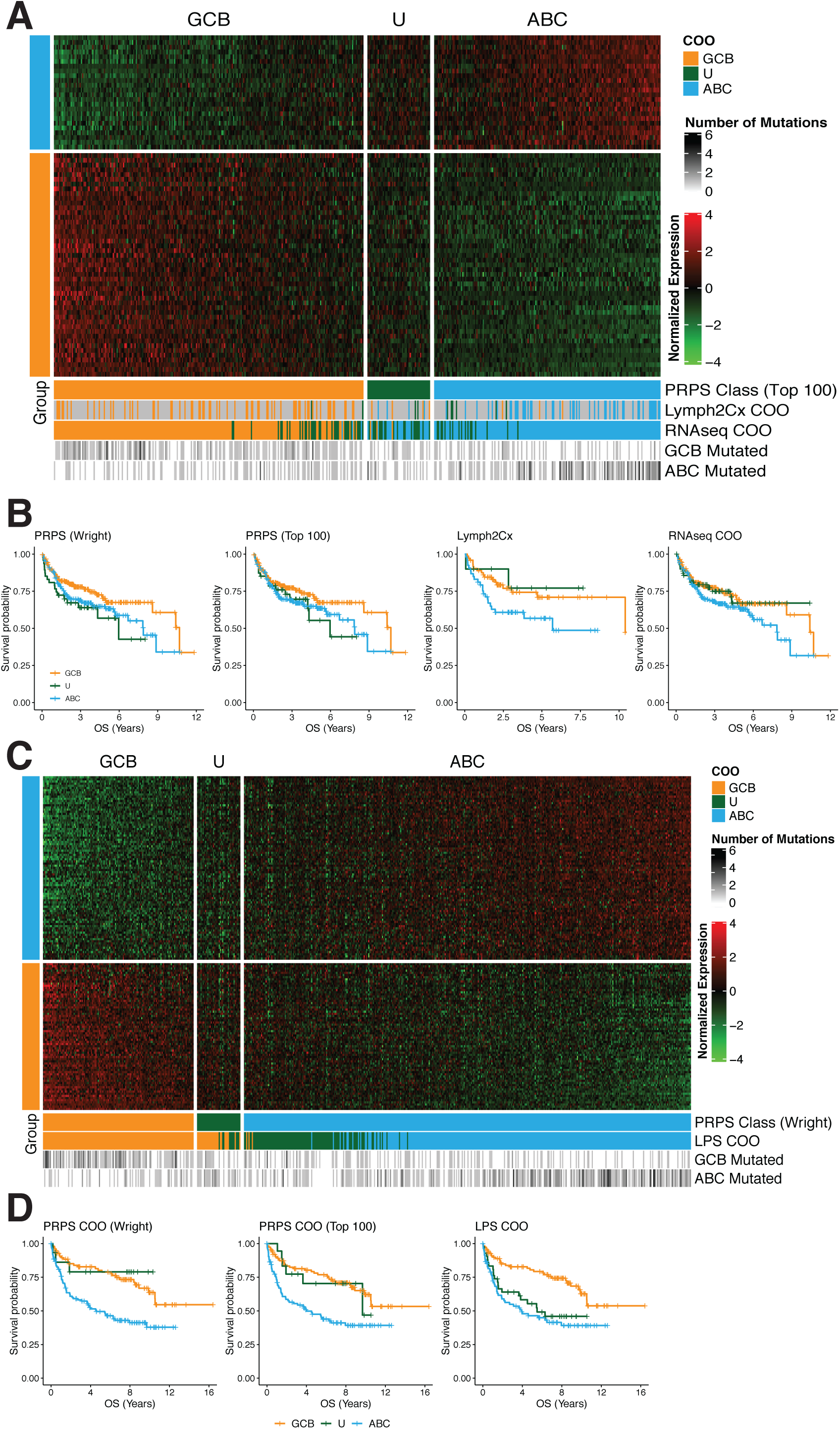
Performance of self-training PRPS COO classification on the Reddy and Schmitz cohorts based on matched top 100 genes and Wright COO genes. **A.** Heatmap of 69 matched “top 100” COO genes from Reddy et al. Samples are ordered by PRPS score. Expression data are normalized per gene to a mean of 0 and standard deviation of 1. **B**. Kaplan–Meier OS (overall survival) survival curves for different COO classifications with Reddy data. **C**. Heatmap of matched 153 Wright COO genes from Schmitz et al. Expression data are normalized per gene to a mean of 0 and standard deviation of 1. **D**. Kaplan–Meier OS (overall survival) survival curves for different COO classifications with Schmitz data.

### Application of self-training to DHITsig sub-classification within GCB

We next sought to validate the PRPS self-training method for sub-classification of GCB-DLBCL using the double hit signature (DHITsig) [15]. This signature was designed to identify DLBCLs with both *MYC* and *BCL2* translocations (genetic double hit) along with tumors having similar biology that may lack one or both genetic features. In contrast to the COO classification shown here, where the ABC and GCB classes are similar in proportions, the DHITsig+ class is generally only present in about 20-40% of GCB-DLBCLs, providing an opportunity to determine accuracy of self-training with imbalanced classes. The REMoDL-B cohort was selected for validation because of its large size and the availability of *MYC* and *BCL2* break apart FISH data for many tumors, providing opportunity for approximating accuracy. Of 543 GCB tumors, 152 (27%) were classified as DHITsig+, and, as expected, all genetic double hit tumors in this cohort were classified as DHITsig+ with PRPS self-training (Figure 4A). Of 98 DHITsig+ tumors with available FISH data, 32 (33%) were *MYC* and *BCL2* double hit. This is a smaller proportion of genetic double hit tumors than we observed in the Ennishi cohort, where ∼ 50% of DHITsig+ tumors are double hit. However, consistent with our initial description of this group, DHITsig+ GCB-DLBCL exhibited inferior PFS and OS relative to DHITsig-GCB-DLBCL (Figure 4B-C). In Cox models adjusted for genetic double hit status, the DHITsig+ class is the only group with significantly inferior outcomes (Table 4). These results are consistent with our observations of DHITsig in other cohorts [15].

**Table 4.**

| Outcome | Covariate | Effect | Total n | Hazard ratio | 95% CI lower | 95% CI upper | P-value |
| --- | --- | --- | --- | --- | --- | --- | --- |
| OS | PRPS Class | INDETERMINATE (vs DHITsig-) | 245 | **1.33** | 0.28 | 6.28 | **0.7165** |
|  |  | DHITsig+ (vs DHITsig-) |  | **4.31** | 1.86 | 9.99 | **0.0007** |
|  | Genetic DHIT | POS (vs NEG) | 245 | **1.60** | 0.76 | 3.36 | **0.2114** |
| PFS | PRPS Class | INDETERMINATE (vs DHITsig-) | 245 | **0.84** | 0.25 | 2.83 | **0.7752** |
|  |  | DHITsig+ (vs DHITsig-) |  | **2.51** | 1.36 | 4.60 | **0.0031** |
|  | Genetic DHIT | POS (vs NEG) | 245 | **1.73** | 0.91 | 3.27 | **0.0926** |

**Figure 4:**
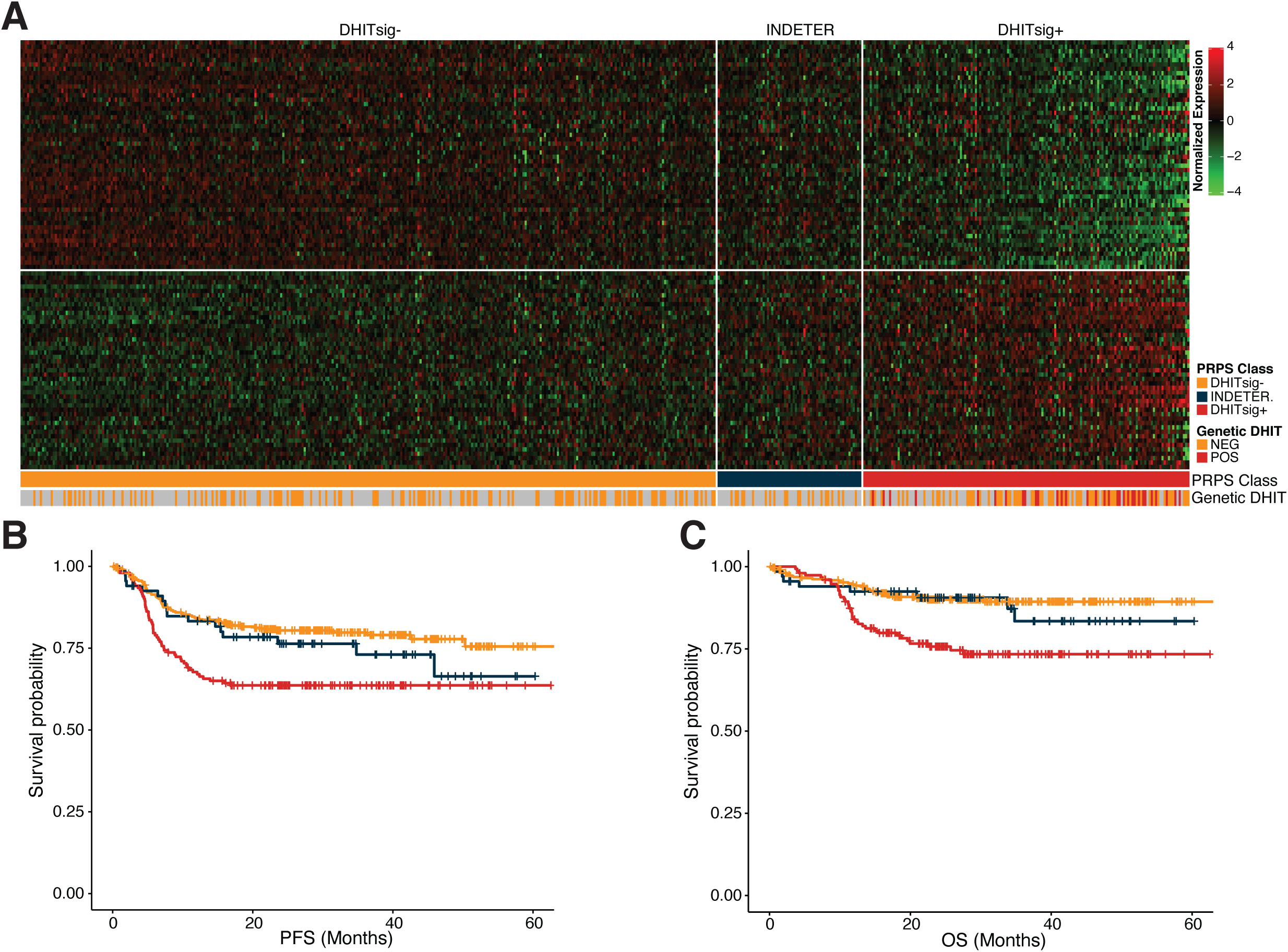
Self-training performance of DHITsig classification on the REMoDL-B GCB cohort. **A**. Heatmap of 97 matched DHITsig genes. Expression data are normalized per gene to a mean of 0 and standard deviation of 1. **B**. Kaplan–Meier PFS (Progression-free survival) and OS (overall survival) survival curves of DHITsig classification.

### Stability of self-training across sample/cohort sizes

In order to establish the relationship between cohort size and self-training classification accuracy, we performed random sub-sampling and COO or DHITsig classification of the Reddy, Scott, and REMoDL-B cohorts. Random sampling and classification were repeated 1000 times for each sample size, and the classification results were compared to the PRPS self-training classifications obtained using the full cohort (Figure 5A). Overall, the samplings of the Scott cohort exhibited higher accuracy than the Reddy cohort for COO classification, which may be because the largest sample size (110) is a larger proportion of the total cohort size (119). Based on the results of the 3×3 accuracy tests, the 25^th^ percentiles are all >= 80% when sample sizes >= 40. Therefore, we recommend that the total sample size should be at least 40 for a balanced binary classification such as COO.

**Figure 5:**
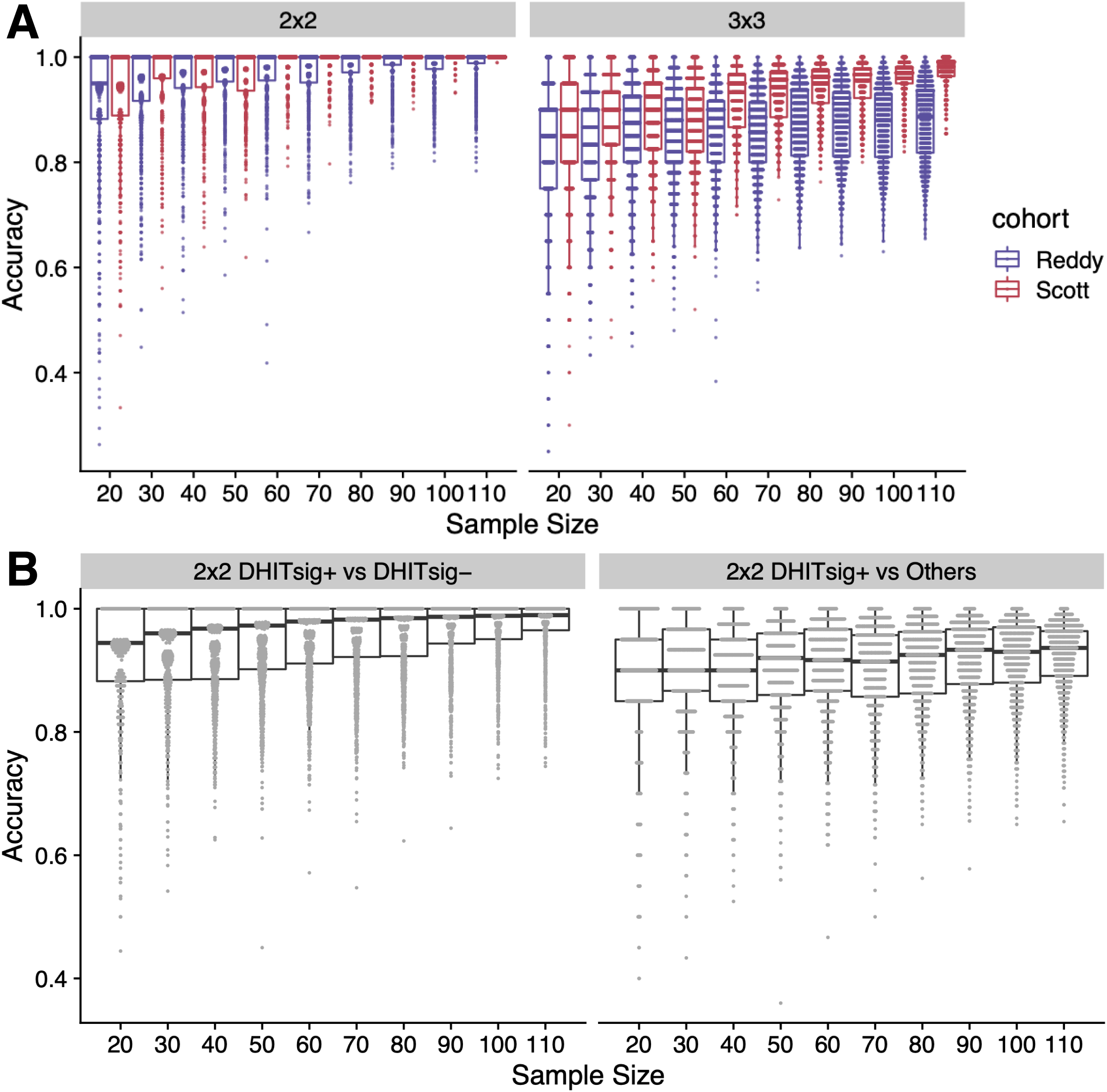
Accuracy distribution of 1000 random samplings for a variety of sample sizes. **A**. COO classification accuracy plots for Reddy and Scott cohorts. **B**. DHITsig classification accuracy plots for the REMoDL-B GCB cohort.

To address the class imbalance in DHITsig classification, the REMoDL-B cohort was sampled to maintain the proportion of DHITsig+ cases (27%) in all tests. We observed that 2-class accuracy with DHITsig+ and DHITsig-classes was greater than 90% across the range of sample sizes, and 2×2 DHITsig+ vs others (DHITsig- and indeterminate) accuracy was greater than 85% across the range (Fig. 5B). Based on the 25^th^ percentile of the 2×2 DHITsig+ vs others accuracy samplings, the sample sizes should not be smaller than 50 in tests with this degree of class imbalance. Larger sample sizes may be needed for classifications with more imbalance between classes.

## Discussion

Traditional supervised classification requires a training data set with ground truth classification information whereby a model is derived from the labeled cases and then used to assign class labels to a testing data set. Importantly, the underlying assumption is that subsequent data sets are comparable to the initial training data. This is often not easy to be satisfied in practice, *e.g.*, in cross-laboratory [32] and cross-platform situations [10,33]. This can limit the reproducibility of discoveries made on different gene expression platforms or in projects involving multiple laboratories. Various normalization methods are often used in an attempt to force compatibility between data sets, however, this introduces a risk of removing true signal and increasing noise, which may result in random or systematic classification errors [10,11]. Numerous classifiers initially developed for application to cancer were implemented using training data from legacy platforms such as microarrays, including the popular COO classification used in DLBCL research [13].

Here, we have evaluated a new binary classifier that can be trained in the absence of comparable labeled training data through self-training. We have demonstrated that gene lists and weights derived from a single training set allow automatic class assignment even on data generated from different laboratories and distinct gene expression platforms. Our algorithm is robust even with incomplete correspondence in genes quantified by different assays, an issue that would require re-training of model-based classifiers such as multivariable regression or SVM. Notably, in the process of porting the 104-gene DHITsig PRPS classification model to the NanoString DLBCL90 assay, we reduced the number of features to 30 without an appreciable decrease in the accuracy of the classifier [15], further supporting the robustness of this method.

Despite the clear importance of the set of genes used in developing a classifier, there is no consistent approach for selecting an optimal set of genes. Here, we applied COO self-training classification using two gene lists: Wright and top 100. All 153 Wright genes were significantly differentially expressed between COO groups respectively in the three cohorts, but many of those unique to this gene set had more modest *t* values (Figure S1). This observation likely relates to differences in gene expression platform used in the process of defining COO. Among both gene sets there is variation in the magnitude of *t* values between RNA-seq cohorts, which also suggests that technical variation can affect the relative importance of these genes. Many of the genes unique to the Wright list thus naturally have a smaller influence on the classifier due to lower weights, such that the results are consistent with either gene list (Figure 2-3, Table 2-3, Figure S3-S5, Supplemental Tables S15-17).

While retaining comparable accuracy to standard methods for COO assignment, our approach consistently left fewer cases unclassified (Figure 2, Figure S3-S4). Relying on the presence of COO-associated mutations, we demonstrated that the mutation status of cases classified as ABC and GCB was consistent with the genetics of these subgroups in both the Reddy and Schmitz cohorts (Figure 6, Table 3). This confirms that the increased classification rate did not cause a decrease in accuracy, and the higher classification rate can be considered another benefit of this method. Similarly, for the REMoDL-B cohort, we assessed the accuracy of DHITsig classification using the presence of genetic features that underlie many DHITsig+ cases. As expected, all tumors with *MYC* and *BCL2* translocations (genetic double hit) were classified into DHITsig+ group (Figure 4).

To obtain sufficient pseudo training data, PRPS self-training requires a sufficiently large sample size and representation of both classes. However, many situations naturally impose limits on the number of samples available. For scenarios in which both classes are naturally represented in approximately equal proportions, our data indicate that a sample size of at least 40 is sufficient for adequate self-training (Figure 5A). Our sampling experiments show that, for situations in which there is a minor class representing ∼20% of cases (DHITsig+), a sample size of 50 is sufficient (Figure 5B). We expect self-training to work in more extreme cases of class imbalance, although this would require a larger sample size.

Taken together, we have shown that our new self-training method is an effective tool for binary classification in the absence of comparable training and testing data sets. Our method represents a significant advance over existing classification methods, addressing many of the caveats usually associated with porting classification models to new data. We expect it to improve the consistency of gene expression-based binary classification across many different cancers.

## Acknowledgements

BC Cancer Centre for Lymphoid Cancer gratefully acknowledges research funding support from the Terry Fox Research Institute (1043), Genome Canada, Genome BC, the Canadian Institutes for Health Research, and the BC Cancer Foundation. RDM holds an ASH Foundation Junior Scholar award and is a Michael Smith Foundation for Health Research Scholar. DWS is supported by a Michael Smith Foundation for Health Research Health Professional-Investigator award and the BC Cancer Foundation. The authors thank George Wright for helpful discussions regarding the Scott cohort. We also gratefully acknowledge the patient donors of samples used herein.

## Availability of data and materials

PRPS is an open-source R package available at GitHub (https://github.com/ajiangsfu/PRPS). PRPS self-training function is “PRPSstableSLwithWeights”.

Gene expression data used to test PRPS self-training algorithm are from Ennishi et al., Reddy et al., Schmitz et al., Scott et al., and Sha et al. [15,16,23,24].

The datasets supporting the conclusions of this article are included within the article and its additional files.

### Contributions

AJ developed and programmed PRPS package including self-training algorithm. JT and CKR generated mutation data sets. BMG and LKH performed pre-processing of the expression data. AJ and LKH performed all classification experiments and generated the figures. AJ, LKH and RDM together wrote the manuscript. RDM directed the project with input from DWS. All authors read and approved the final manuscript.

## Ethics declaration

This project was approved by the BC Cancer research ethics board.

## References

1. Heo YJ, Park C, Yu D, Lee J, Kim K-M. Reproduction of molecular subtypes of gastric adenocarcinoma by transcriptome sequencing of archival tissue. Sci Rep. 2019;9:1–8.

2. Solin LJ, Gray R, Baehner FL, Butler SM, Hughes LL, Yoshizawa C, et al. A multigene expression assay to predict local recurrence risk for ductal carcinoma in situ of the breast. J Natl Cancer Inst. 2013;105:701–10.

3. Kopetz S, Tabernero J, Rosenberg R, Jiang Z-Q, Moreno V, Bachleitner-Hofmann T, et al. Genomic classifier ColoPrint predicts recurrence in stage II colorectal cancer patients more accurately than clinical factors. Oncologist. 2015;20:127–33.

4. Golub TR, Slonim DK, Tamayo P, Huard C, Gaasenbeek M, Mesirov JP, et al. Molecular Classification of Cancer: Class Discovery and Class Prediction by Gene Expression Monitoring. Science. 1999;286:531–7.

5. Huang S, Cai N, Pacheco PP, Narrandes S, Wang Y, Xu W. Applications of Support Vector Machine (SVM) Learning in Cancer Genomics. Cancer Genomics Proteomics. 2018;15:41–51.

6. Algamal ZY, Alhamzawi R, Mohammad Ali HT. Gene selection for microarray gene expression classification using Bayesian Lasso quantile regression. Comput Biol Med. 2018;97:145–52.

7. Toh K-A, Lin Z, Sun L, Li Z. Stretchy binary classification. Neural Netw. 2018;97:74–91.

8. Ayyad SM, Saleh AI, Labib LM. Gene expression cancer classification using modified K-Nearest Neighbors technique. Biosystems. 2019;176:41–51.

9. Podolsky MD, Barchuk AA, Kuznetcov VI, Gusarova NF, Gaidukov VS, Tarakanov SA. Evaluation of Machine Learning Algorithm Utilization for Lung Cancer Classification Based on Gene Expression Levels. Asian Pac J Cancer Prev. 2016;17:835–8.

10. Zhang S, Shao J, Yu D, Qiu X, Zhang J. MatchMixeR: a cross-platform normalization method for gene expression data integration. Bioinformatics. 2020: https://doi.org/10.1093/bioinformatics/btz974

11. Vu T, Riekeberg E, Qiu Y, Powers R. Comparing normalization methods and the impact of noise. Metabolomics. 2018;14:108.

12. Lenz G, Wright GW, Emre NCT, Kohlhammer H, Dave SS, Davis RE, et al. Molecular subtypes of diffuse large B-cell lymphoma arise by distinct genetic pathways. 2008;105:13520–13525.

13. Wright G, Tan B, Rosenwald A, Hurt EH, Wiestner A, Staudt LM. A gene expression-based method to diagnose clinically distinct subgroups of diffuse large B cell lymphoma. Proc Natl Acad Sci. 2003;100:9991–6.

14. Alizadeh AA, Eisen MB, Davis RE, Ma C, Lossos IS, Rosenwald A, et al. Distinct types of diffuse large B-cell lymphoma identified by gene expression profiling. Nature. 2000;403:503–511.

15. Ennishi D, Jiang A, Boyle M, Collinge B, Grande BM, Ben-Neriah S, et al. Double-Hit Gene Expression Signature Defines a Distinct Subgroup of Germinal Center B-Cell-Like Diffuse Large B-Cell Lymphoma. JCO. 2018;37:190–201.

16. Scott DW, Wright GW, Williams PM, Lih C-J, Walsh W, Jaffe ES, et al. Determining cell-of-origin subtypes of diffuse large B-cell lymphoma using gene expression in formalin-fixed paraffin-embedded tissue. Blood. 2014;123:1214–1217.

17. Scrucca L, Fop M, Murphy TB, Raftery AE. mclust 5: Clustering, Classification and Density Estimation Using Gaussian Finite Mixture Models. R J. 2016;8:289–317.

18. Liao Y, Smyth GK, Shi W. featureCounts: an efficient general purpose program for assigning sequence reads to genomic features. Bioinformatics. 2014;30:923–30.

19. Love MI, Huber W, Anders S. Moderated estimation of fold change and dispersion for RNA-seq data with DESeq2. Genome Biol. 2014;15:550.

20. Dobin A, Davis CA, Schlesinger F, Drenkow J, Zaleski C, Jha S, et al. STAR: ultrafast universal RNA-seq aligner. Bioinformatics. 2013;29:15–21.

21. Morin RD, Mendez-Lago M, Mungall AJ, Goya R, Mungall KL, Corbett RD, et al. Frequent mutation of histone-modifying genes in non-Hodgkin lymphoma. Nature. 2011;476:298–303.

22. Reddy A, Zhang J, Davis NS, Moffitt AB, Love CL, Waldrop A, et al. Genetic and Functional Drivers of Diffuse Large B Cell Lymphoma. Cell. 2017;171:481–494.e15.

23. Schmitz R, Wright GW, Huang DW, Johnson CA, Phelan JD, Wang JQ, et al. Genetics and Pathogenesis of Diffuse Large B-Cell Lymphoma. N Engl J Med. 2018;378:1396–1407.

24. Sha C, Barrans S, Cucco F, Bentley MA, Care MA, Cummin T, et al. Molecular High-Grade B-Cell Lymphoma: Defining a Poor-Risk Group That Requires Different Approaches to Therapy. JCO. 2018;37:202–12.

25. Patro R, Duggal G, Love MI, Irizarry RA, Kingsford C. Salmon provides fast and bias-aware quantification of transcript expression. Nat Methods. 2017;14:417–9.

26. Anders S, Pyl PT, Huber W. HTSeq--a Python framework to work with high-throughput sequencing data. Bioinformatics. 2015;31:166–9.

27. Kim S, Scheffler K, Halpern AL, Bekritsky MA, Noh E, Källberg M, et al. Strelka2: fast and accurate calling of germline and somatic variants. Nat Methods. 2018;15:591–4.

28. Chen X, Schulz-Trieglaff O, Shaw R, Barnes B, Schlesinger F, Källberg M, et al. Manta: rapid detection of structural variants and indels for germline and cancer sequencing applications. Bioinformatics. 2016;32:1220–2.

29. Karczewski KJ, Francioli LC, Tiao G, Cummings BB, Alföldi J, Wang Q, et al. The mutational constraint spectrum quantified from variation in 141,456 humans. bioRxiv. Cold Spring Harbor Laboratory; 2020;531210.

30. Zhao H, Sun Z, Wang J, Huang H, Kocher J-P, Wang L. CrossMap: a versatile tool for coordinate conversion between genome assemblies. Bioinformatics. 2014;30:1006–7.

31. Arthur SE, Jiang A, Grande BM, Alcaide M, Cojocaru R, Rushton CK, et al. Genome-wide discovery of somatic regulatory variants in diffuse large B-cell lymphoma. Nat Commun. 2018;9:4001.

32. Doyle RM, O’Sullivan DM, Aller SD, Bruchmann S, Clark T, Coello Pelegrin A, et al. Discordant bioinformatic predictions of antimicrobial resistance from whole-genome sequencing data of bacterial isolates: an inter-laboratory study. Microb Genom. 2020;6.

33. Xu X, Zhang Y, Williams J, Antoniou E, McCombie WR, Wu S, et al. Parallel comparison of Illumina RNA-Seq and Affymetrix microarray platforms on transcriptomic profiles generated from 5-aza-deoxy-cytidine treated HT-29 colon cancer cells and simulated datasets. BMC Bioinformatics. 2013;14 Suppl 9:S1.

