## Supplemental Figures for "Robust gene expression-based classification of cancers without normalization"

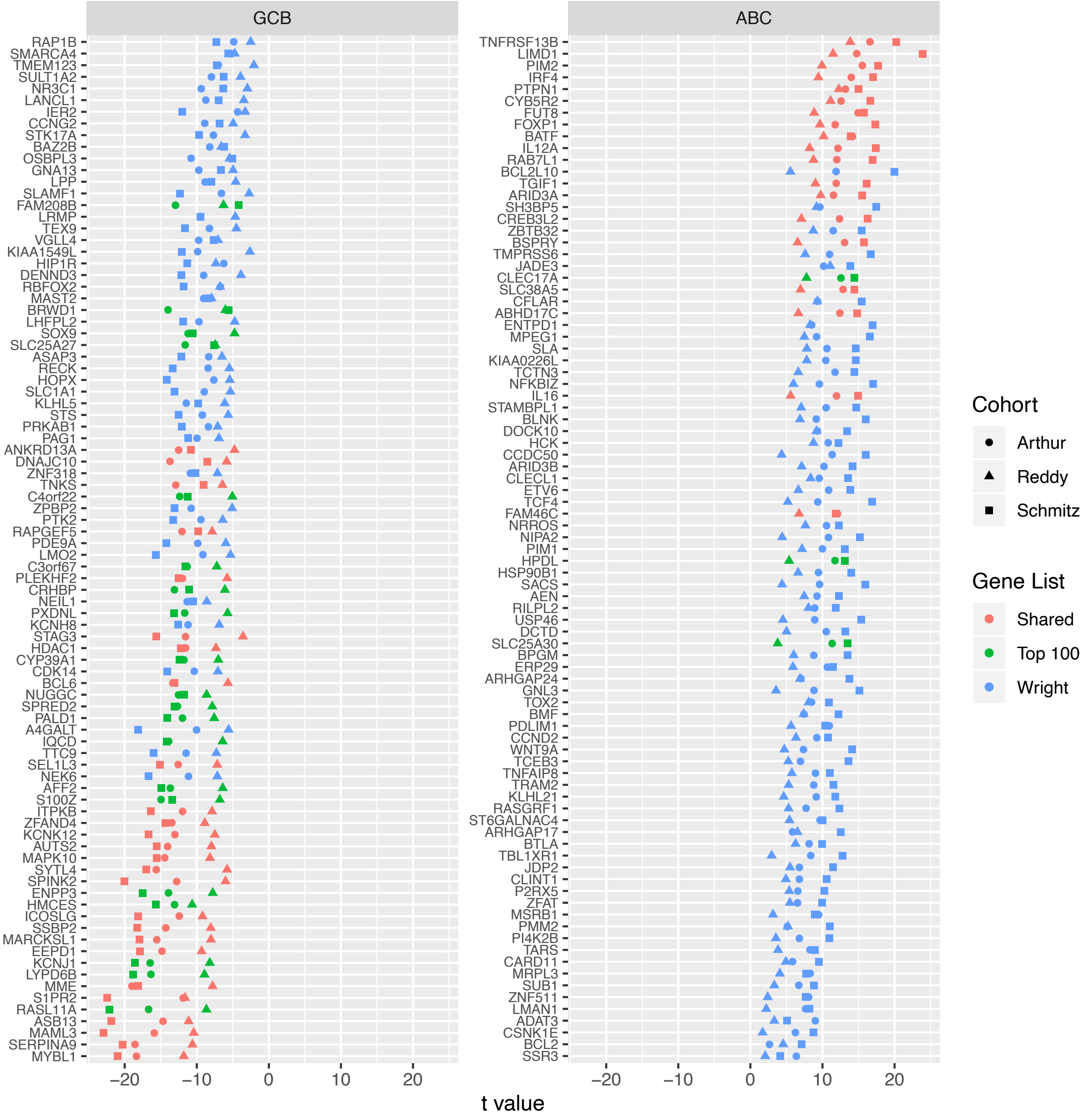


**Figure S1: Comparison of *t* values of Wright and top 100 gene sets used for COO classification in the Ennishi, Reddy, and Schmitz cohorts.**

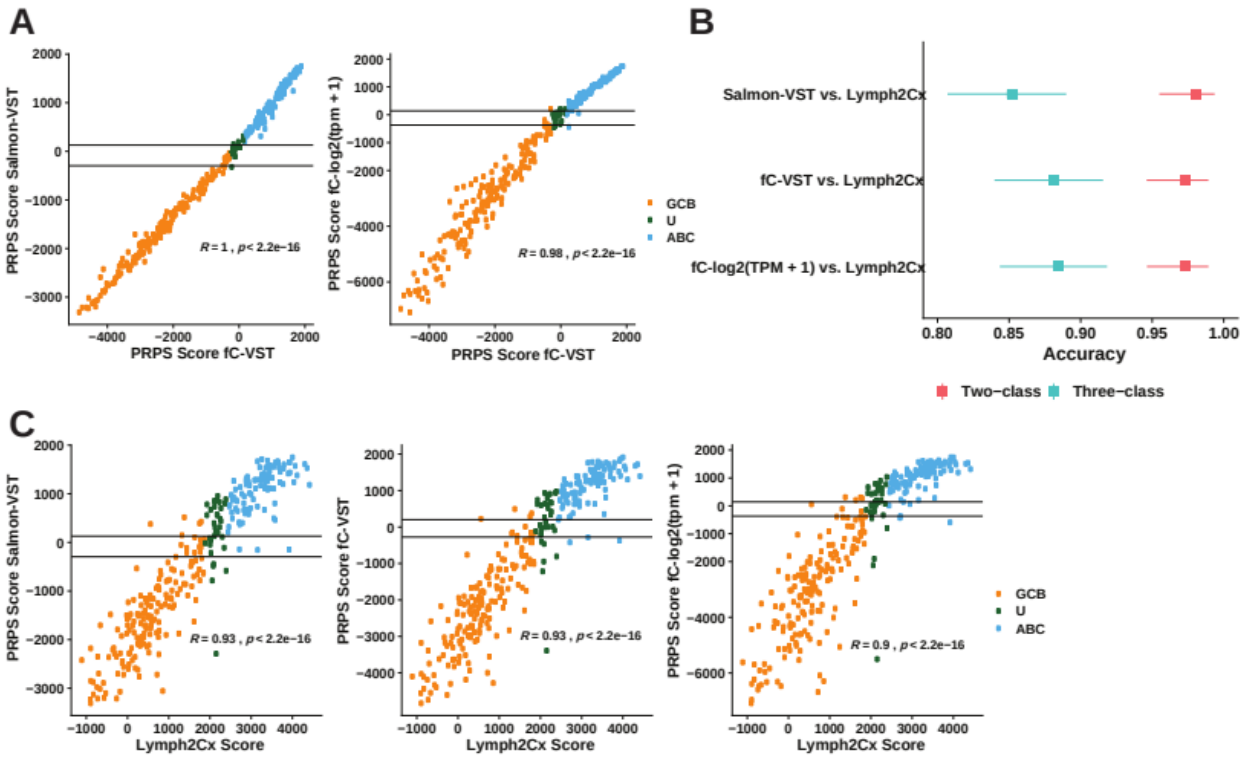


**Figure S2: Self-learning performance of COO classification on different expression data formats with Ennishi cohort. A**. COO classification score plots comparing results from three data formats. **B**. Accuracy comparison plot. 95% confidence intervals are in shown bars. **C**. COO classification score plots comparing results of three data formats to Lymph2Cx.

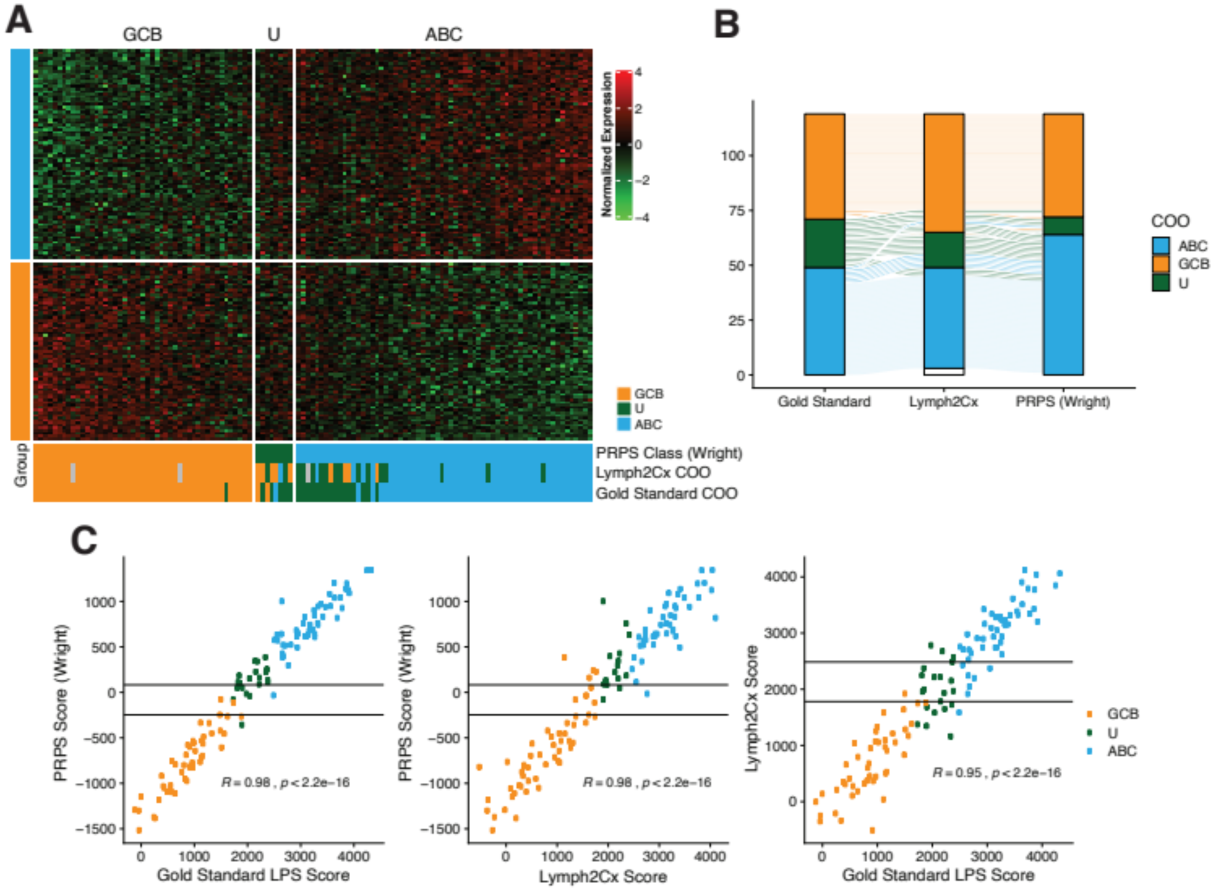


**Figure S3: Self-learning performance of COO classification on the Scott cohort with matched 151 Wright COO genes. A**. Heatmap of matched 151 Wright genes. The expression data are normalized/standardized to mean as 0 and standard deviation as 1 for each gene. **B**. COO classification migration plot. **C**. COO classification score plots.

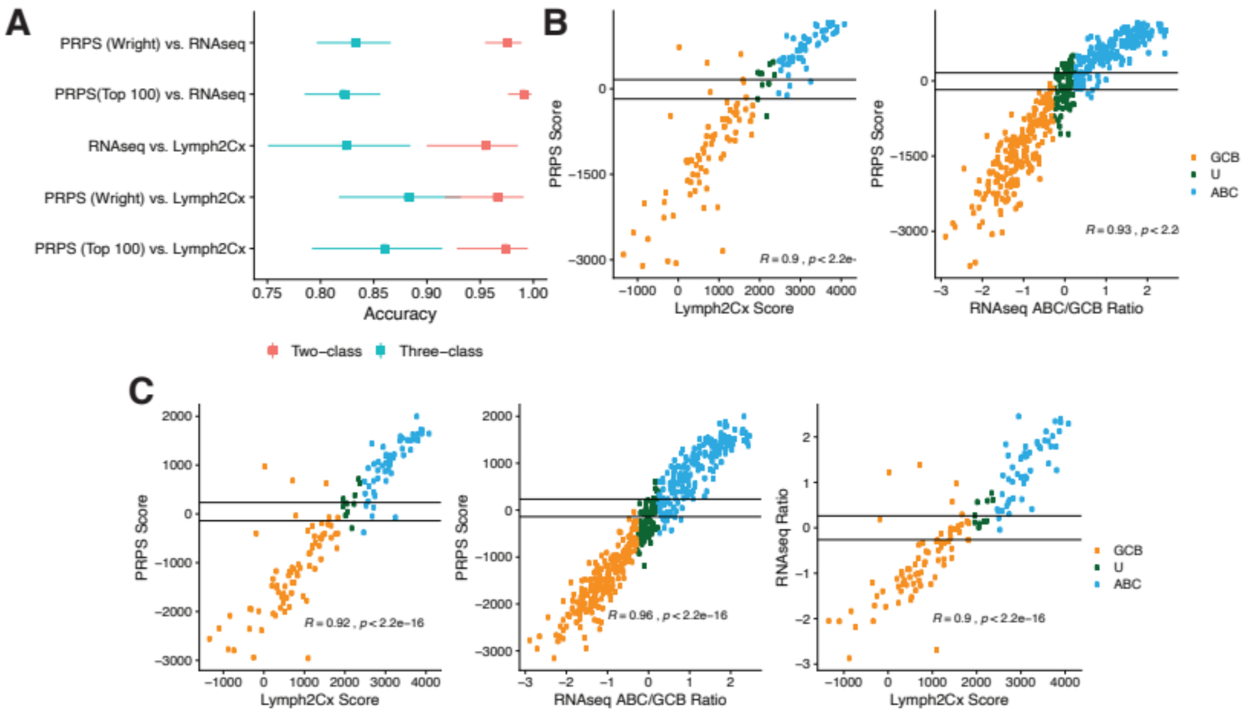


**Figure S4: Self-learning performance of COO classification on the Reddy cohorts based on matched top 100 COO genes developed from Ennishi cohort and matched Wright COO genes.** **A**. Accuracy comparison plot. 95% confidence intervals are shown in bars. **B**. COO classification score plots of PRPS with matched 71 top 100 genes. **C**. COO classification score plots of PRPS with matched 152 Wright genes and more.

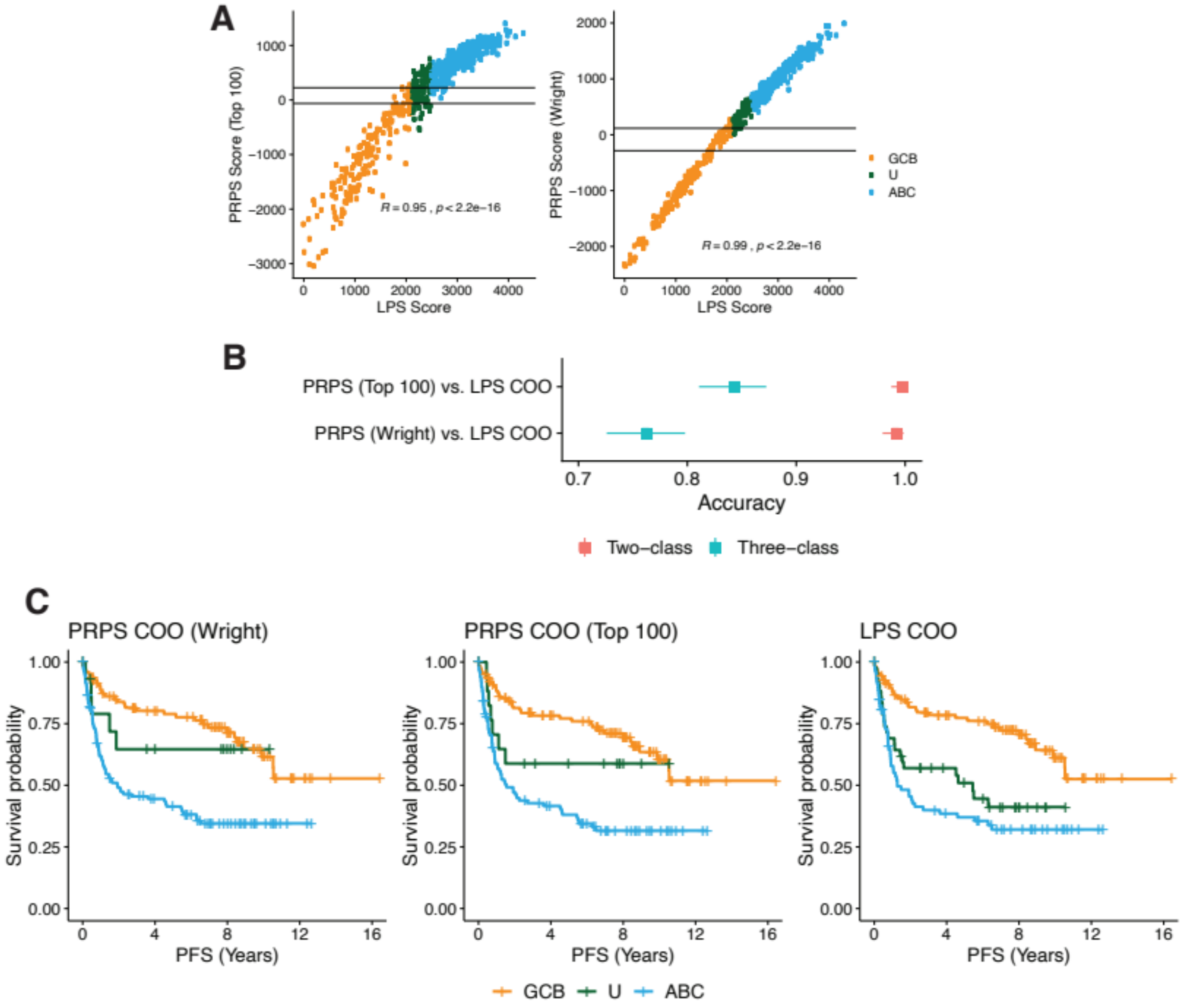


**Figure S5: Self-learning performance of COO classification on the Schmitz cohorts based on matched top 100 COO genes developed from Ennishi cohort and Wright COO genes. A**. COO classification score plots. **B**. Accuracy comparison plot. Pink for two-class accuracy. 95% confidence intervals are shown in bars. **C**. Kaplan–Meier PFS (Progression-free survival) survival curves of COO classification.

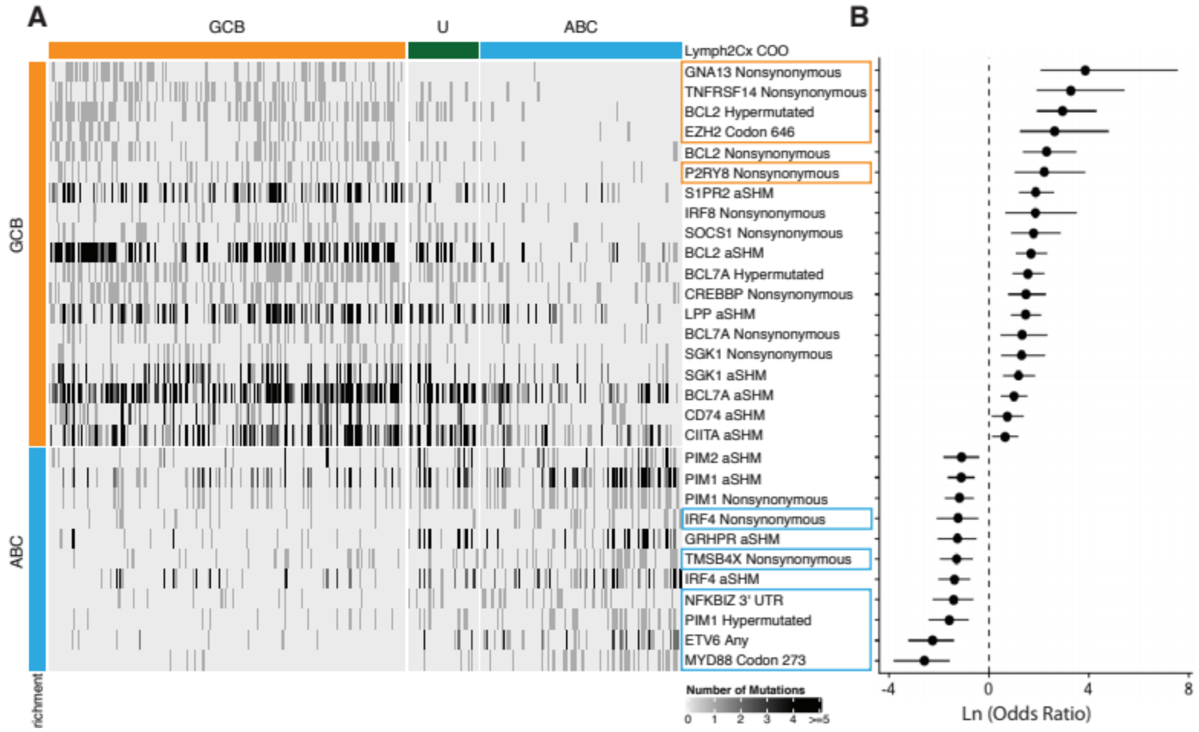


**Figure S6: COO mutation enriched genes from Ennishi cohort.** **A**. Mutation incidence and frequency for COO-enriched genes. The number of mutations affecting for genes known to be aSHM targets includes any mutation affecting an exon. For NFKBIZ, mutations in the 3' UTR (a mutation hot spot) are shown. For other genes, only non-synonymous mutations are shown. **B**. Odds ratio forest plot with values transformed with natural log (ln).
